# β-Caryophyllene Enhances the Transcriptional Upregulation of Cholesterol Biosynthesis in Breast Cancer Cells

**DOI:** 10.1101/569558

**Authors:** Mam Y. Mboge, Andrea Ramirez-Mata, Adam Bullock, Riley O’Donnell, John V. Mathias, Julie Davila, Christopher J. Frost, Susan C. Frost

## Abstract

β-caryophyllene (BCP) exhibits anti-proliferative properties in cancer cells. Here, we examine the hypothesis that BCP induces membrane remodeling. Our data show that high concentrations of BCP increase membrane permeability of human breast cells (hBrC) causing detachment and cell death. At a sub-lethal concentration of BCP, we show that BCP induces a striking upregulation of genes involved in cholesterol biosynthesis, including the gene that encodes for HMGCoA reductase (HMGCR), the rate-determining step in cholesterol biosynthesis. In addition, stearoyl-CoA desaturase (SCD) is also upregulated which would lead to the enhanced formation of monounsaturated fatty acids, specifically oleate and palmitoleate from stearoyl CoA and palmitoyl CoA, respectively. These fatty acids are major components of membrane phospholipids and cholesterol esters. Together, these data suggest that cells respond to BCP by increasing the synthesis of components found in membranes. These responses could be viewed as a repair mechanism and/or as a mechanism to mount resistance to the cytotoxic effect of BCP. Blocking HMGCR activity enhances the cytotoxicity of BCP, suggesting that BCP may provide an additional therapeutic tool in controlling breast cancer cell growth.

## Introduction

World-wide, breast cancer is the most common form of cancer in women (1). Failure to treat metastatic disease is the leading cause of death (2). Although substantial progress has been made towards treating cancer, such as surgical resections or adjuvant therapies, highly aggressive cancers remain a challenge. One of the deadliest types of cancers in women is triple negative breast cancer (TNBC), where expression of the estrogen, progesterone, and HER2 receptors is either low or absent (3). This type of cancer is one of the most difficult to treat due to lack of targeted therapies and its inherent therapeutic resistance. TNBC accounts for a disproportionate number of breast cancer deaths (4).

Notable success in the search for novel anti-cancer drugs in natural products has been achieved from the unique chemical reactions associated with plants. Plants create remarkably complex chemicals beyond those that are required for their own growth and reproduction (5). For generations, these “specialized” metabolites have been used in the treatment of human diseases, even though their active components and molecular targets are not always well-defined (6). While specialized metabolites are derived from a number of biosynthetic pathways, terpenes are the most abundant class of plant secondary metabolites (7). In plants, terpenes are synthesized through the traditional mevalonate pathway (MVP) which produces the intermediates, isopentenyl pyrophosphate and dimethylallyl pyrophosphate from which terpenes derive, but also via the methylerythritol phosphate pathway (8). The MVP produces an array of monoterpenes, diterpenes, tetraterpenes, and precursors of complex sterols (9,10). Some plant terpenoids are already well-established for the treatment of breast cancer. For example, paclitaxel (sold under the trade name of Taxol) is an oxygenated diterpenoid that was originally isolated from the Pacific Yew and first shown to stabilize microtubules in the Horwitz lab (11) As noted by Heinig et al., synthetic paclitaxel is arguably the most successful anticancer drug of all time (12), and is one of the cytotoxic drugs of choice for the treatment of triple negative and drug resistant (recurrent) breast cancers (13).

Over the last decade, a number of investigators have demonstrated that β-caryophyllene (BCP) exhibits anti-proliferative properties in cancer cells (6,14–19). BCP is a bicyclic sesquiterpene (Figure 1) and a major plant volatile of many plant essential oils (like cloves, oregano, black pepper, cinnamon, and cannabis). BCP is often found with small quantities of isocaryophyllene, the oxide form of caryophyllene, and humulene. It was the first known “dietary cannabinoid” and has achieved GRAS (Generally Recognized as Safe) status. Because of its unique taste and pleasant odor, BCP has been used in cosmetics and as a flavoring since the mid 1900’s and has FDA approval. It also has well-known anti-pathogenic properties in plants (20), as well as anti-inflammatory (17,21), anti-nociceptive (22), and anti-mutagenic (23) properties in animals. Over a decade ago, Legault and Pichette demonstrated that BCP stimulates the accumulation of paclitaxel in DLD-1 (colon cancer) cells, implicating enhanced membrane permeability as a mechanism of action (19). Indeed, BCP has been shown to interact directly with phospholipid bilayers increasing membrane fluidity (24). Earlier studies in *E.coli* showed that cyclic hydrocarbons, including terpenes, interact directly with biological membranes (25). Accumulation of these hydrocarbons results not only in increased membrane fluidity, but also in membrane swelling, both signs of cell stress. At biological temperatures, membrane fluidity is controlled by the saturation state of the acyl chains of fatty acids (primarily in phospholipids) and cholesterol content (26). Changes in either of these parameters could lead to membrane remodeling which can affect membrane function.

**Figure 1.**
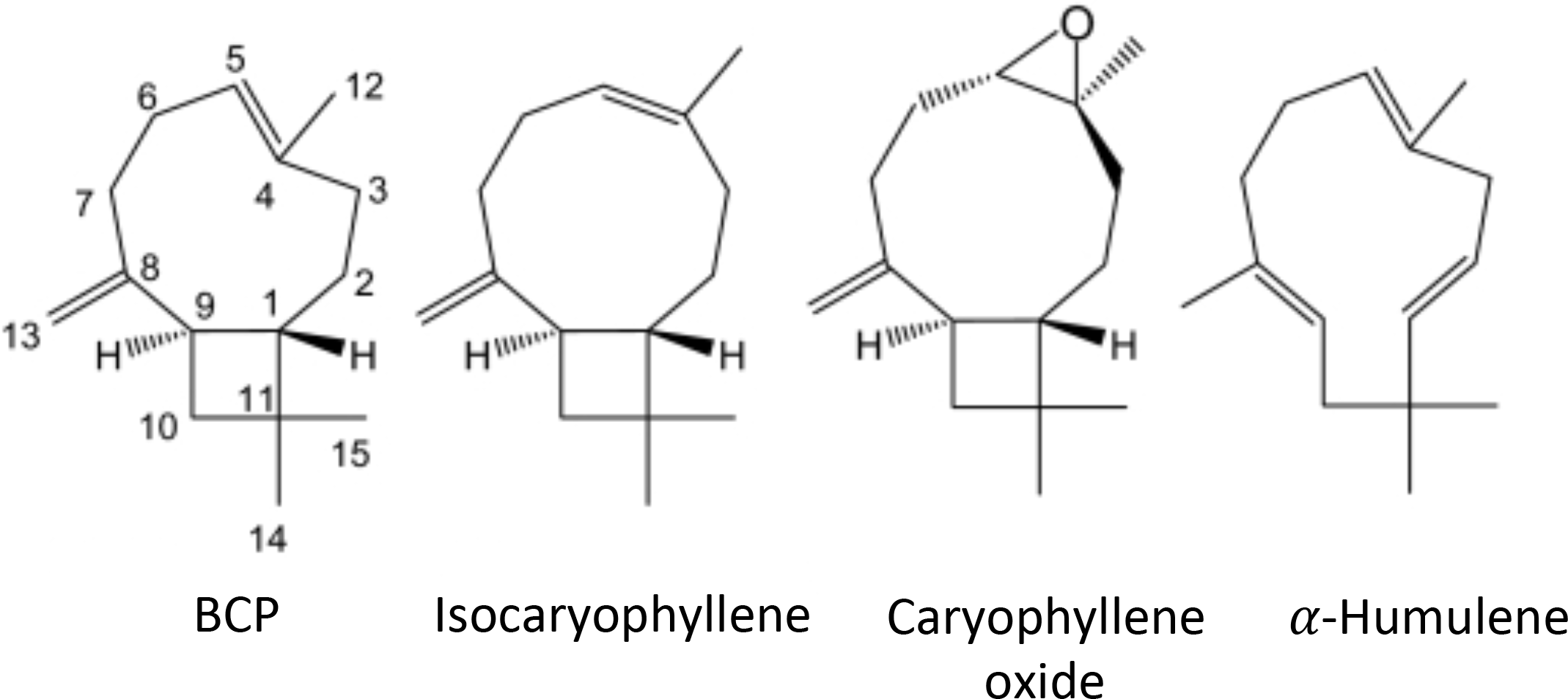
Isomeric structures of caryophyllene. β-Caryophyllene (BCP) accounts for ~98-99% found in nature (65). Other forms of caryophyllene are also shown in this figure that exist with BCP but are less abundant. BCP is a strong agonist for the cannabinoid receptor 2 (CB_2_). This figure is adapted from that in Gertsch *et al*. (16).

Here, we examine the hypothesis that BCP induces membrane remodeling. Our data show that high concentrations of BCP increase membrane permeability of human breast cells (hBrC) causing detachment and cell death. At a “sub-lethal” concentration of BCP, we show that BCP induces a striking upregulation of genes involved in cholesterol biosynthesis, including the gene that encodes for HMGCoA reductase (HMGCR), the rate-determining step in cholesterol biosynthesis. Many of the identified genes are regulated by the transcription factor SREBP-2 whose activation is regulated by cholesterol availability (for review see (27)). In addition, stearoyl-CoA desaturase (SCD1) is also upregulated. This endoplasmic reticulum (ER) enzyme catalyzes the rate-limiting step in the formation of monounsaturated fatty acids, specifically oleate and palmitoleate from stearoyl CoA and palmitoyl CoA, respectively.

These fatty acids are major components of membrane phospholipids, cholesterol esters, and alkyl-diacyglycerol. Together, these data suggest that cells respond to sub lethal concentrations of BCP by increasing the synthesis of components found in membranes. These responses could be viewed both as a repair mechanism and potentially as a mechanism to mount resistance to the cytotoxic effect of BCP. We hypothesize that blocking the repair mechanism would enhance the cytotoxicity of BCP. This might provide an addition therapeutic tool in controlling breast cancer cell growth assuming that specificity could be established.

## Materials and Methods

### Cell Culture

All cell lines were authenticated. MCF10A cells were a gift from Dr. Brian Law. Dr. Keith Robertson provided T47D cells. The MDA-MB-231-LM2 cells were a gift from Dr. Dietmar Siemann and the UFH-001 line was developed in the lab of Dr. Susan Frost (28), both of which have the triple negative phenotype. Each line was maintained at 37°C at 5% CO_2_. MCF10A cells were cultivated in Dulbecco’s Modified Eagle’s Medium (DMEM/Ham’s F12 medium (1:1) (Corning Lellgro) supplemented with 5% horse serum (Sigma Aldrich), 10 μg/ml insulin (Eli Lilly), 20 ng/mL epidermal growth factor (EGF) (Upstate Biochem) and 100 ng/mL dexamethasone (Sigma Aldrich). The T47D cells were maintained in McCoy’s medium (Gibco) containing 10% fetal bovine serum (FBS; Sigma Aldrich) and 1μg/mL bovine insulin (Elanco). The MDA-MB-231-LM2 and UFH-001 cell lines were cultivated in DMEM supplemented with 10% FBS.

### Viability assays

To measure the viability of cells exposed to β-caryophyllene (Sigma/Aldrich # 22075), we measured mitochondrial function using thiazolyl blue (MTT) from Sigma/Aldrich (# M5655). Briefly, cells were seeded in 96 well plates and treated with increasing concentration of drugs. The cells were incubated with BCP for 4h, 8 h, and 24 h under normoxic conditions. In some experiments, we tested the effect of paclitaxel for a duration of 24 h or 48 h. Thiazolyl blue was added 4 h before the end of each time point. The medium was then removed without disrupting the cells and DMSO was added for 15 min to each well. Absorbance was read at 570 nm using an Epoch microplate reader (Biotek). Data represent the average ± SEM of three biological replicates.

### Cytotoxicity assays

To measure the cytotoxic effects of BCP and paclitaxel, we used the release of lactate dehydrogenase (LDH), measured as the production of NAD (Sigma/Aldrich, # MA0K066). Cells were grown in 35 mm dishes and exposed to specific concentrations of BCP at times indicated in the figure legends. In some experiments, we tested the effect of paclitaxel over 48 h. At high concentrations of BCP, but not of paclitaxel, it was noted that cells were released from the plate. Medium was separated from cells by centrifugation (13,200xg for 5 min) before sampling the medium for LDH activity. Absorbance was read at 450 nm using the Epoch microplate reader. LDH activity is reported as nmoles/min/10^6^ cells, based on cell counts at the start of each experiment. Data represent the mean ± SEM from three 3 biological replicates.

### RNA isolation and RNAseq

To measure the effect of BCP on global gene expression, we used RNAseq technology. RNA (RIN >9) was extracted from UFH-001 cells, previously exposed to normoxia or hypoxia for 16h in the presence or absence of BCP at either 20 μM or 200 μM (RNase easy plus mini kit from Qiagen). Libraries were prepared at the Genomics Core at the University of Louisville and sequencing was performed on triplicate biological replicates (Illunima NestSeq 500). This generated over 144 million 75 bp reads that aligned to the human genome (96.3% alignment rate), or approximately 24 million reads per sample. The data discussed in this publication have been deposited in NCBI’s Gene Expression Omnibus (29) and are accessible through GEO Series accession number GSE125511 (https://www.ncbi.nih.gov/geo/query/acc.cgi?acc=GSE125511).

### Membrane preparation

Cells were washed 3 times with ice cold PBS. Cells were then homogenized in buffer containing 20 mM Tris base, 1 mM EDTA, and 255 mM sucrose, pH 7.4, supplemented with 1 mM phenylmethyl sulfonyl fluoride (PMSF) and protease inhibitor cocktail (Sigma/Aldrich, P-8340). Membranes were collected by centrifugation at 212,000 × g for 60 min at 4°C. Membranes were resuspended in a small volume of the same buffer and stored at −20°C.

### Lysate preparation

Cells were washed 3 times with ice cold PBS and then extracted in RIPA buffer [1% Triton X-100, 10 mM Tris-HCl (pH 8.0), 140 mM NaCl, 1 mM EDTA, 0.5 mM EGTA, 0.1% sodium deoxycholate, 0.1% SDS] supplemented with PMSF and protease inhibitor cocktail for 15 min on ice. Lysates were clarified by centrifugation at 212,000 × g for 60 min at 4°C. Clarified supernatants were collected and stored at −20°C. Protein concentrations for both membranes and cell lysates was determined using the Markwell modification of the Lowry procedure (30).

### SDS-PAGE and Western blotting

One-dimensional SDS-polyacrylamide gel electrophoresis (SDS-PAGE) was performed essentially as described by Laemmli et al. (31). Protein samples were mixed with sample dilution buffer. Gels (10%) were typically run overnight at room temperature at approximately 45 V in a Hoefer SE 600 electrophoresis unit. Protein samples were electro-transferred from SDS-PAGE gels to nitrocellulose membranes in transfer buffer (25 mM Tris-base, 192 mM glycine, 20% methanol) at 200 mA for 2 h at 4°C. Western blotting was accomplished as previously described (32). Enhanced chemiluminescence (ECL) was used according to manufacturer’s directions (GE Healthcare, #RPN2106 or RPN2232). Santa Cruz antibodies were used for analysis of HMGCR (sc-271595) and Na^+^K^+^ATPase (sc-28800). The GAPDH antibody was from Cell Signaling (D16H11). Band intensity was quantified using Un-Scan-It (Silk Scientific, Inc.) in the linear range of the film.

### Cholesterol Quantification

Cholesterol concentration was estimated using a kit from Sigma/Aldrich (MAK043). Briefly, cells (in 35mm plates) were washed with PBS, and collected by centrifugation in 0.5 mL PBS. After removing the buffer, cholesterol was extracted with 200 μL of chloroform/isopropanol/IGEPAL (ratio of 7:11:0.1) according to instructions by the manufacturer. This was mixed vigorously and then exposed to centrifugal force (13,000 × g) in the cold for 10 min. The organic phase was transfer to a new tub and dried at 50°C. Residual solvent was removed under vacuum (Savant). Dried samples were responded in assay buffer, and mixed on a vortex. This was used to measure total cholesterol. We also assessed the effect of BCP on steady-state uptake of [^3^H]-cholesterol. Cells were equilibrated for 8 h in DMEM containing 10% FBS, spiked with 1uCi/uL [1,2^3^H(N)]cholesterol (NET19001MC, 44.5Ci/mmol). Cells were then exposed to BCP under normoxic and hypoxic conditions for 16 h. Samples were taken from the medium before washing cells with ice-cold PBS. Cells were extracted in 1.0 mL of 0.1% sodium dodecyl sulfate. Samples were collected for counting and protein analysis. Data represent the avg ± SEM of three biological replicates.

### Statistical evaluation

Unless otherwise noted, each experiment was repeated three times and reported as the average ± S.E.M. Statistical analysis was performed using Prism 7 software using the Student’s t-test. *p* values < 0.05 was considered statistically significant and are reported in the Figure legends.

## Results

### Effect of BCP on metabolic function and cytotoxicity in breast cancer cells

Because it has been reported that BCP enhances growth inhibition in a prostate cell line induced by paclitaxel (19), we analyzed the effect of paclitaxel across a panel of human breast cancer cell lines (hBrC). We first determined the effect of paclitaxel on cell growth (Supplementary Figure 1). This panel represents a control line (MCF10A), an ER+ luminal line (T47D), and two triple negative lines (MDA-MB-231-LM2 and UFH-001 (33)). The characteristics of the new UFH-001 line have been described elsewhere (28). We used the MTT assay, which measures mitochondrial function, as a surrogate measure of cell number. At both 24 and 48 h, the MDA-LM2 and UFH-001 lines were more sensitive to growth inhibition by paclitaxel than either the MCF10A and T47D cells (Supplementary Figures 1A and 1B). This was not surprising given that these cells replicate more quickly, in our hands, than the MCF10A or T47D cells. However, we were unable to detect any amplification by 70 μM BCP on paclitaxel induced cell growth (Supplementary Figure 1C). The BCP concentration used here was similar to that used by Legault and Picchet (19). While paclitaxel induced some cytotoxicity in the UFH-001 line as measured by lactate dehydrogenase (LDH) release from cells, BCP at 70 **μ**M did not further influence cytotoxicity of either the UFH cells or the other cells tested (Supplementary Figure 1D). Reasons for these differences between our data and those previously published could be related to the sensitivity of the breast cancer cell lines relative to the prostate line used previously. That said, we must still conclude from our data that paclitaxel action in the cell lines that we tested is not potentiated by BCP.

Earlier studies reported that BCP exhibits anti-cancer activity but that the EC_50_ values vary with cell type (15). We repeated these experiments to determine if BCP is cytotoxic to the four hBrC lines and to establish dose-response curves. We again measured LDH release, this time over 4 h, in the presence or absence of specific concentrations of BCP concentrations. In Figure 1, we show the total LDH activity released during that time frame (Panel A), the activity with the background subtracted (Panel B), and normalized data in percent released (Panel C). In agreement with earlier studies, the sensitivity to BCP varied 2-3 fold across cell lines (Panel C), although the EC_50_ values for all hBrC lines were in high μM concentrations. Concentrations at which we observed elevated LDH activity corresponded to an increase in the number of floating cells. Together, these data demonstrate that BCP interacts with hBrC cells causing detachment from the extracellular matrix, enhancing membrane permeability, and leading to cell death, not necessarily in that order. It is noteworthy that loss of mitochondrial function in UFH cells (Supplementary Figure 1C) occurs at 200 **μ**M while cytotoxicity requires higher concentrations (Figure 1C). This opened an opportunity to investigate mechanisms that might be responsible for BCP action in the absence of cytotoxicity.

### Effect of BCP on transcriptional activity in UFH-001 cells

The UFH-001 line exhibits the TNBC phenotype, are fast growing in culture, and form tumors in mouse models (28). Even though these were not the most sensitive to BCP, we chose to analyze UFH-001cells because targeted therapies for TNBC patients are not available. Because hypoxia is the environment associated with aggressive breast cancers, and an independent prognosticator for poor patient outcome (34), we also chose the condition of hypoxia to evaluate the effect of BCP. In addition, we extended the exposure time from 4 h (Figure 2) to 16 h (Figure 3) to allow for transcriptional responses. We have previously shown that the levels of the hypoxia-inducible transcription factor 1α (HIF1α) increase at the protein level in these cells response to 16 h exposure to both hypoxia and the hypoxic mimic, desferroxamine mesylate (35) confirming their sensitivity to low oxygen conditions. Thus, the cells were exposed of to 1% oxygen (hypoxia) in the presence of 20 **μ**M, 200 **μ**M BCP or vehicle alone (DMSO) over 16 h (Figure 3). We note that there is no difference in BCP-induced cytotoxicity between normoxic and hypoxic cells at 16 h (Supplementary Figure 2).

**Figure 2.**
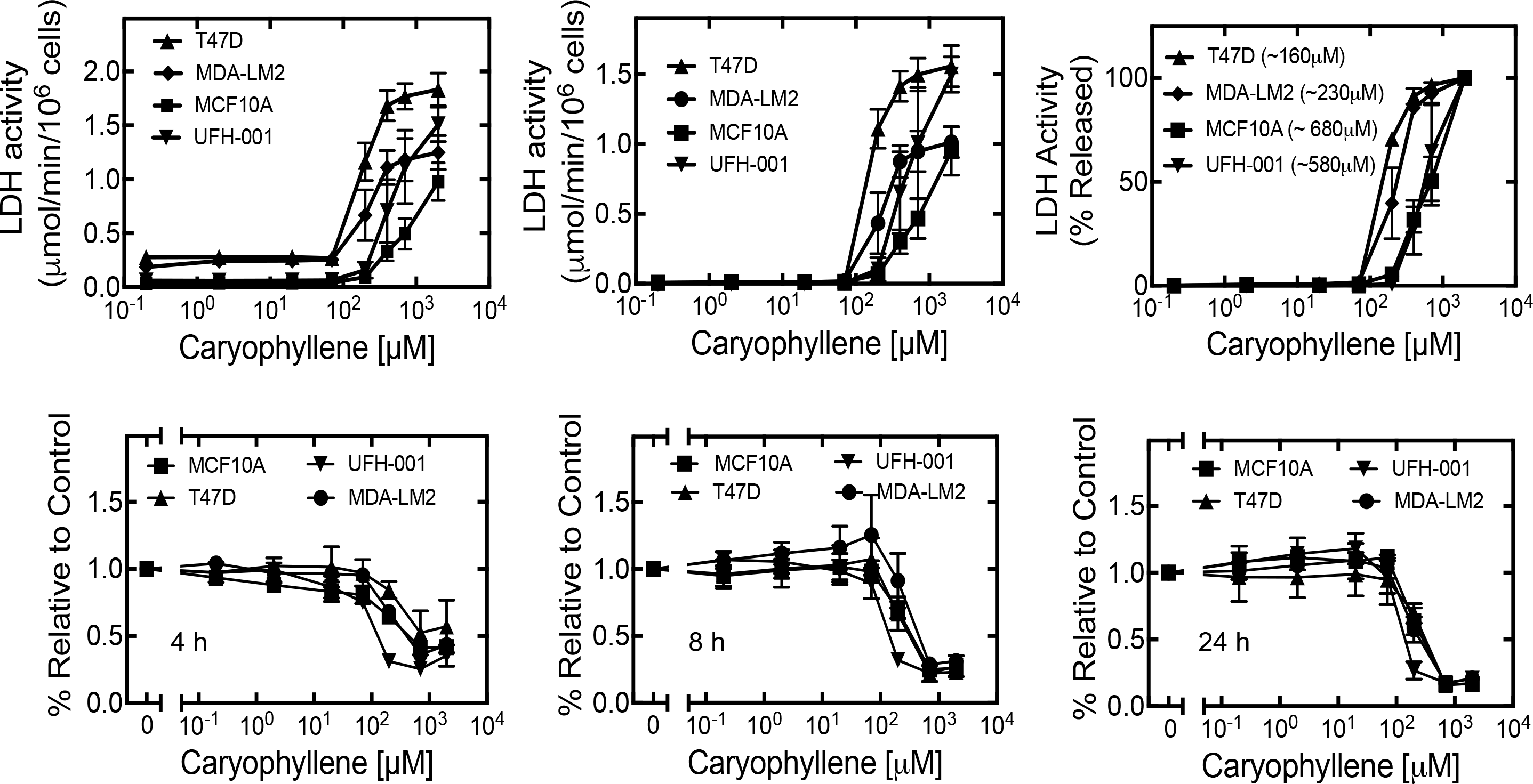
BCP reduces cell viability of breast cancer cells. Cells were treated with increasing concentrations of BCP. LDH release was measured after 4h (**Panels A-C**), showing total activity, background subtracted activity and percent of LDH release relative to the control for each cell line. MTT assays were performed after 4, 8 and 24 h (**Panels D-F**). Data shown represent the average of at least 3 biological replicates ± S.E.M.

**Figure 3.**
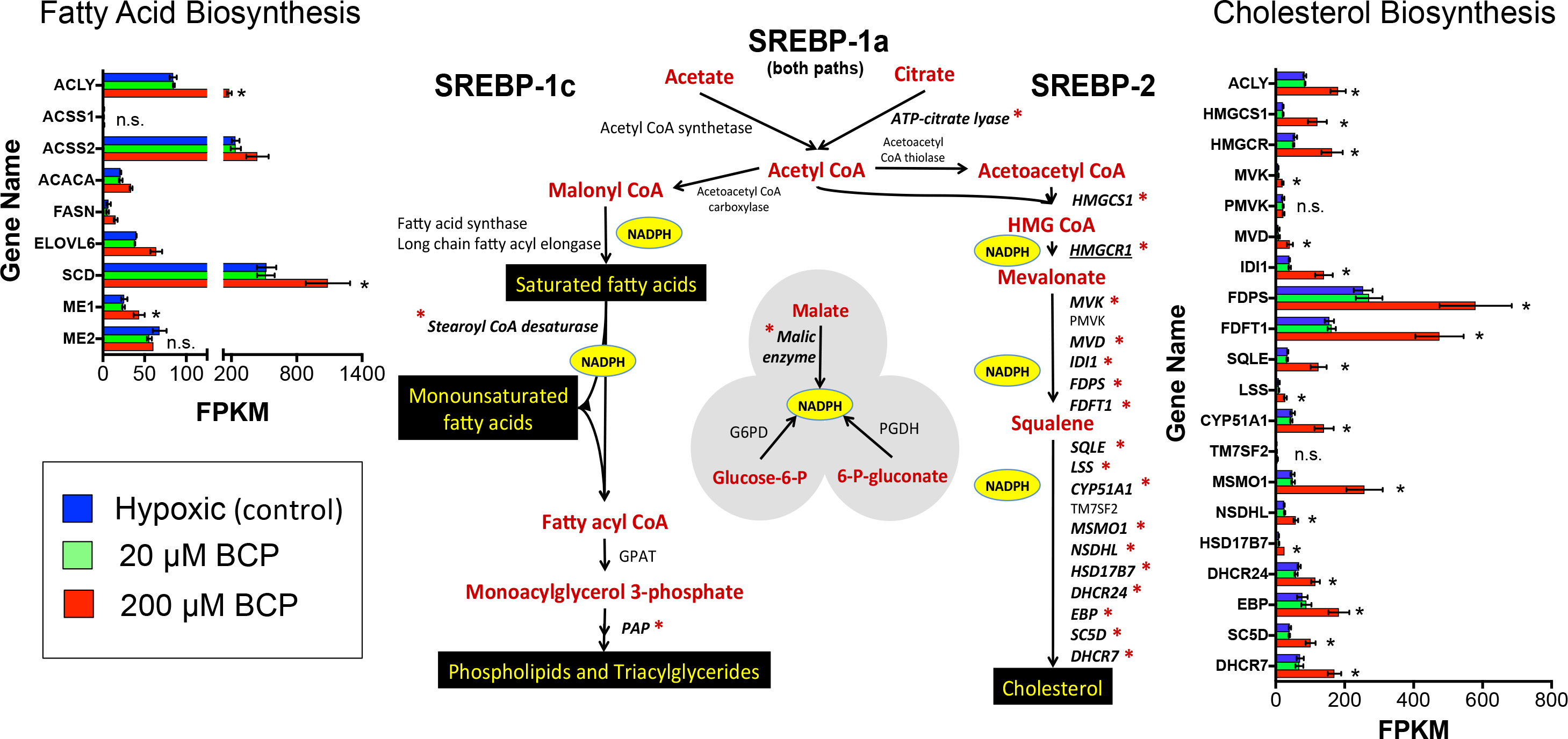
BCP transcriptionally upregulates the genes that affect cholesterol and lipid biosynthesis. RNA was isolated from UFH-001 cells exposed to hypoxia in the absence or presence of BCP. RNAseq was performed at the University of Louisville (see Table 1). Red asterisks next to each gene represent a significant induction (*q* = 0.05) for that gene. Up-regulated genes include those in both the cholesterol and lipid biosynthesis pathways. The central figure was modeled after that in Horton *et al*. (35).

**Table 1.**
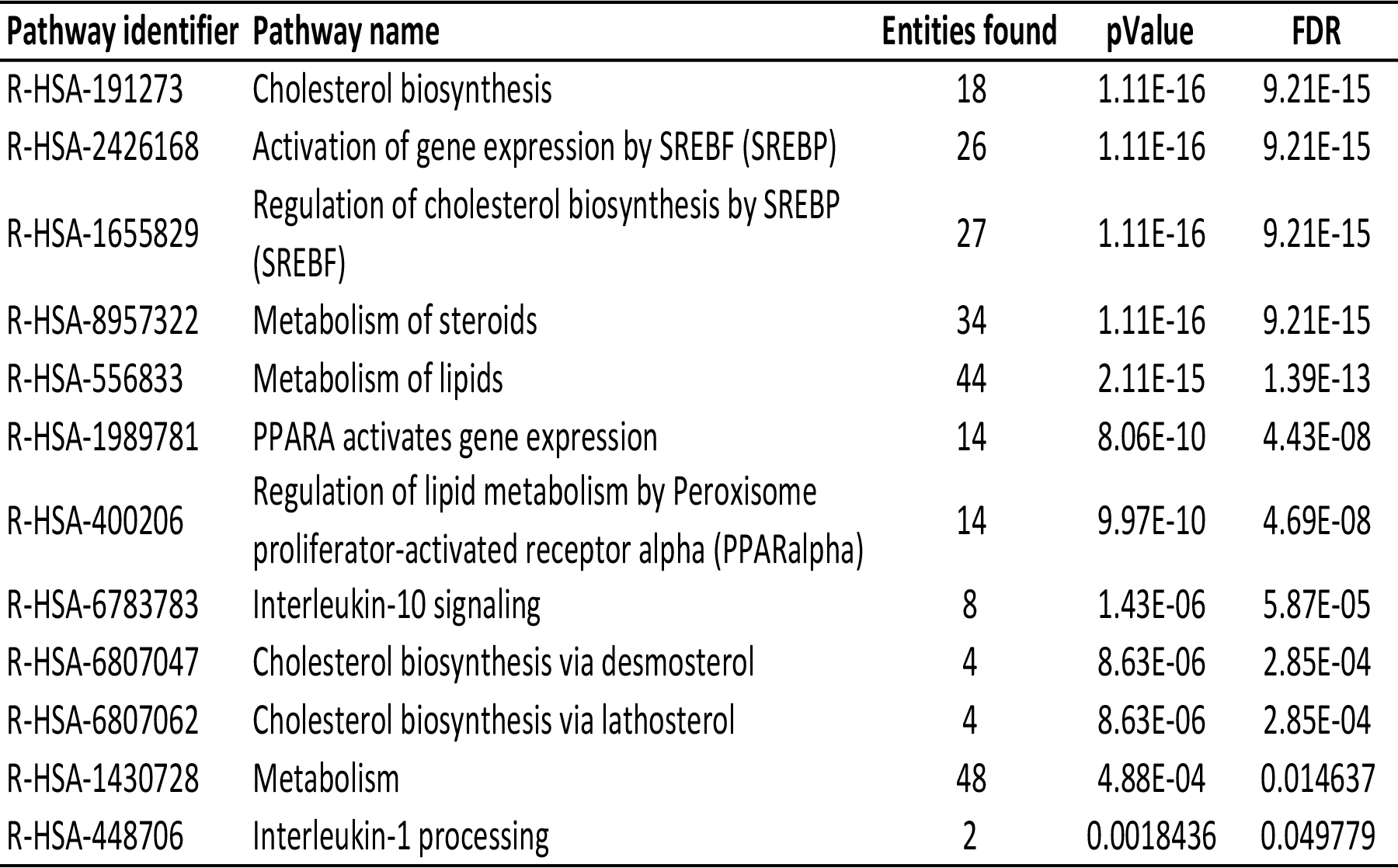
Pathways induced by BCP (RNA-seq pilot data using UFH-001 cells) High quality RNA (RIN < 9) was isolated from UFH-001 cells exposed to hypoxia with and without 200 BCP. The Genomics Core (University of Louisville) prepared libraries and performed sequencing (Illumina NextSeq 500). We selected differentially expressed genes based on FDR adjusted p-values (q-values) ≤ 0.05, and analyzed this gene set for pathway involvement using the Reactome database. Because Reactome also assesses the FDR for pathway analysis, we again selected an FDR <0.05 to enhance specificity.

| Pathway identifier | Pathway name | Entities found | pValue | FDR |
| --- | --- | --- | --- | --- |
| R-HSA-191273 | Cholesterol biosynthesis | 18 | 1.11E-16 | 9.21E-15 |
| R-HSA-2426168 | Activation of gene expression by SREBF (SREBP) | 26 | 1.11E-16 | 9.21E-15 |
| R-HSA-1655829 | Regulation of cholesterol biosynthesis by SREBP (SREBF) | 27 | 1.11E-16 | 9.21E-15 |
| R-HSA-8957322 | Metabolism of steroids | 34 | 1.11E-16 | 9.21E-15 |
| R-HSA-556833 | Metabolism of lipids | 44 | 2.11E-15 | 1.39E-13 |
| R-HSA-1989781 | PPARA activates gene expression | 14 | 8.06E-10 | 4.43E-08 |
| R-HSA-400206 | Regulation of lipid metabolism by Peroxisome proliferator-activated receptor alpha (PPARalpha) | 14 | 9.97E-10 | 4.69E-08 |
| R-HSA-6783783 | Interleukin-10 signaling | 8 | 1.43E-06 | 5.87E-05 |
| R-HSA-6807047 | Cholesterol biosynthesis via desmosterol | 4 | 8.63E-06 | 2.85E-04 |
| R-HSA-6807062 | Cholesterol biosynthesis via lathosterol | 4 | 8.63E-06 | 2.85E-04 |
| R-HSA-1430728 | Metabolism | 48 | 4.88E-04 | 0.014637 |
| R-HSA-448706 | Interleukin-1 processing | 2 | 0.0018436 | 0.049779 |

RNA was isolated from cells to determine changes in the transcriptome of cells exposed to BCP. As a first approach, we selected differentially expressed genes based on FDR adjusted p-values (q-values) of <0.05, and analyzed this gene set for pathway involvement using the Reactome database. Because Reactome also assesses the FDR for pathway analysis, we again selected an FDR <0.05 to enhance specificity. Data complied in Table 1 show that lipid metabolism in pathways regulated by the SREBP family of transcription factors are key targets of BCP action. Also of note in Table 1 is the activation of inflammatory processes through interleukin 10 (IL10) signaling and processing of interleukin 1 (IL1) suggesting that the cells are experiencing ROS-mediated stress.

The exceptional specificity of BCP action is illustrated in Figure 3, modeled in part after Horton et al. (36). The asterisks indicate those genes that are significantly induced by BCP (q ≤ 0.05). Remarkably, 17 of the 19 genes in cholesterol biosynthesis (including HMGCR, the rate-limiting step in the pathway) are induced by BCP. Interestingly, elevated expression of this gene is associated with poor prognosis in TNBC, but not estrogen receptor-positive breast cancer patients (Figure 4) according to the Kaplan-Meier data base (kmplot.com/analysis) using a web tool developed by Lanczky *et al*. to query the databases (37). The coordinated regulation of cholesterol biosynthesis in the UFH-001 cells is reminiscent of those revealed by the early studies of Brown and Goldstein (38). Moreover, our results suggest that BCP modulates fatty acid saturation, as stearoyl CoA desaturase (SCD1) is also upregulated (Figure 3). SCD1 is a transmembrane, ER-resident protein whose function is to create monosaturated fatty acids, specifically oleate and palmitoleate from stearoyl CoA and palmitoyl CoA, respectively. These are major components of membrane phospholipids, and suggest that BCP-treated cells may respond to membrane damage by enhancing the biosynthetic path for both fatty acids and cholesterol.

**Figure 4.**
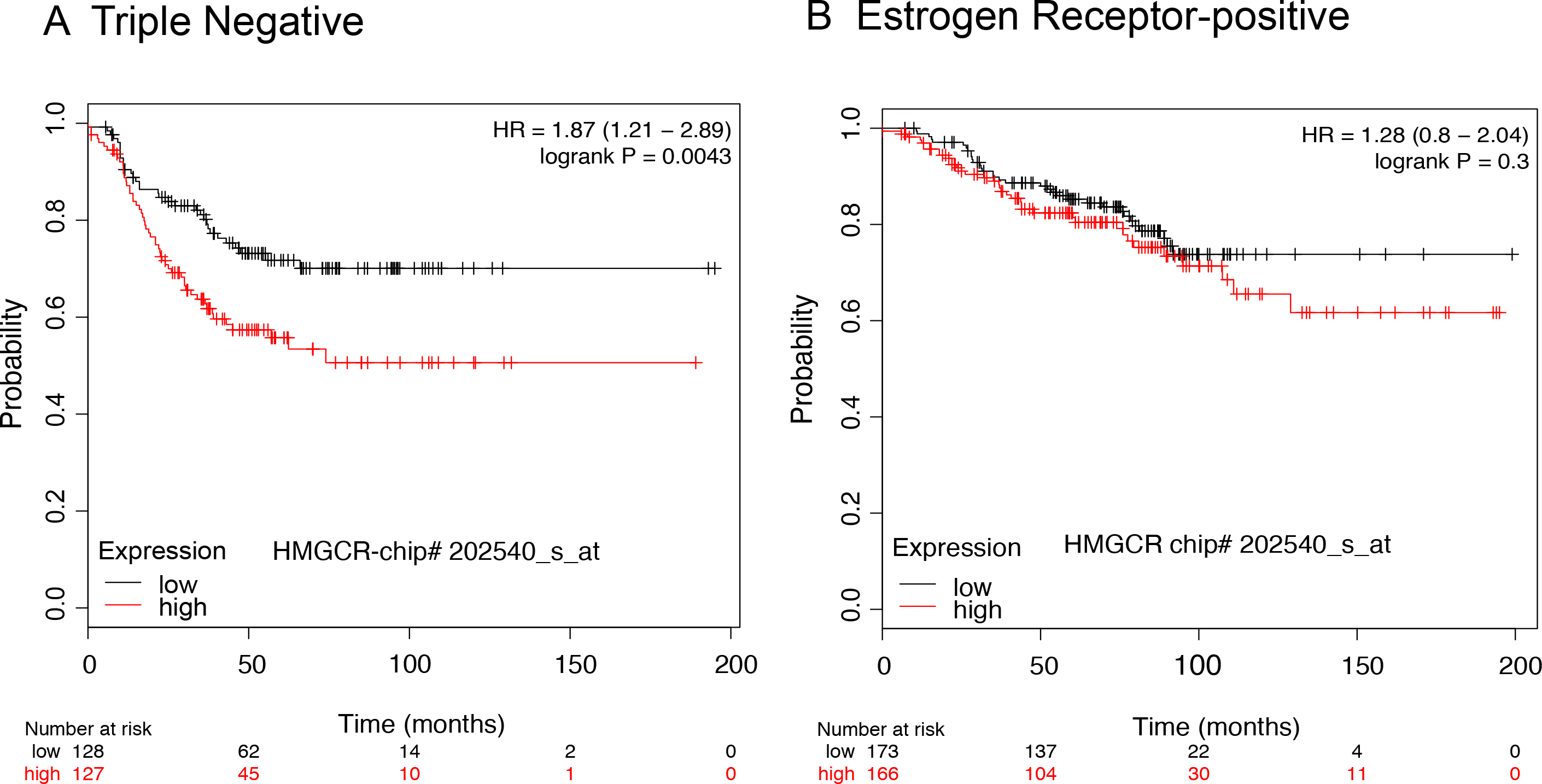
HMGCoA Reductase is a poor prognosticator for triple negative breast cancer patient survival. Kaplan Meier survival curves are shown in Panels A and B. **Panel A** represents survival curves (RFS) of TNBC patients with low or high expression of HMGCoA reductase (*p* = 0.0043). **Panel B** represents survival curves (RFS) of ER-positive patients with low or high expression of HMGCoA reductase (no significance). These data were compiled using a web tool designed by Lanczky *et al*. (36) to analyze data in the Kaplan Meier database

### BCP increases HMGCR expression and cholesterol content

Western blot analysis was used to assess the expression of the HMGCR. BCP had no affect on HMGCR expression under normoxic conditions (Figures 5B,C), although expression was higher than under hypoxic conditions. In contrast, HMGCR expression was increased significantly by 200 **μ**M relative to vehicle control (Figures 5B,C) replicating the RNAseq data (Figures 3 and 5A). This correlated with the accumulation of cholesterol (Figure 5D) where only hypoxic treatment in the presence of 200 μM achieved significance relative to vehicle control. We also measured the ability of cells to accumulate external [^3^H]cholesterol, assuming that this may represent association within the plasma membrane. Surprisingly, significant accumulation was detected in the presence of 200 μM BCP under both normoxic and hypoxic conditions (Figure 5E).

**Figure 5.**
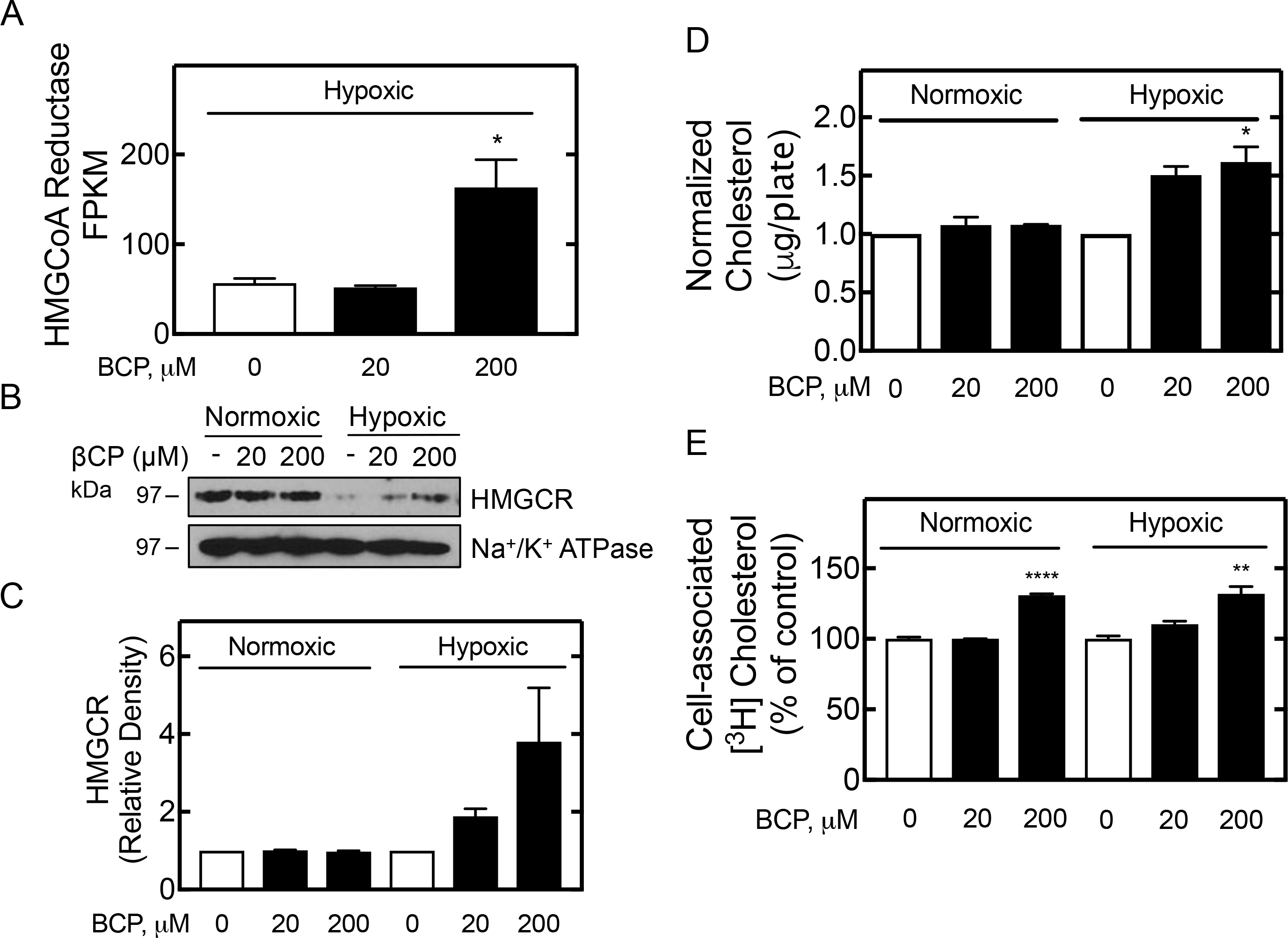
BCP regulates HMGCoA reductase expression in response to BCP in UFH-001 cells. **Panel A.** FPKM values for HMG CoA reductase, n = 3. * q = 0.013. **Panel B.** Membranes were isolated from UFH-001 cells exposed or not to hypoxia in the presence or absence of BCP for 24 h. HMGCoA reductase expression was assessed by western blotting. UN-SCAN-IT (vs 5.3) was used to quantify of HMG CoA reductase expression in three biological replicates ± S.E.M. **Panel C**. Analysis of bands (UN-SCAN-IT vs 5.3) from replicate western blots of HMG CoA Reductase in Panel A are presented as relative density. **Panel D**. Total cholesterol was quantified after Folch extraction of UFH-001 cells, treated as in Panel A, using the Cholesterol Quantification Kit from Sigma/Aldrich (MAK043). These data represent three biological replicates and are reported as μg/μL extract ± S.E.M., relative to DMSO controls (*, p = 0.038). **Panel E**. Radioactive cholesterol associated with UFH-001 cells was measured after preloading cells for 8 h with tracer amounts of [^3^H]cholesterol (~ 480,000 cpm/mL medium) followed by exposure to normoxia or hypoxia, with or without BCP as in Panel A. Data are reported as percent of normoxic or hypoxic controls, based on cpm/μg protein, in triplicate samples ± S.E.M. **, p < 0.01: ****, p = 0.0001.

**Figure 6.**
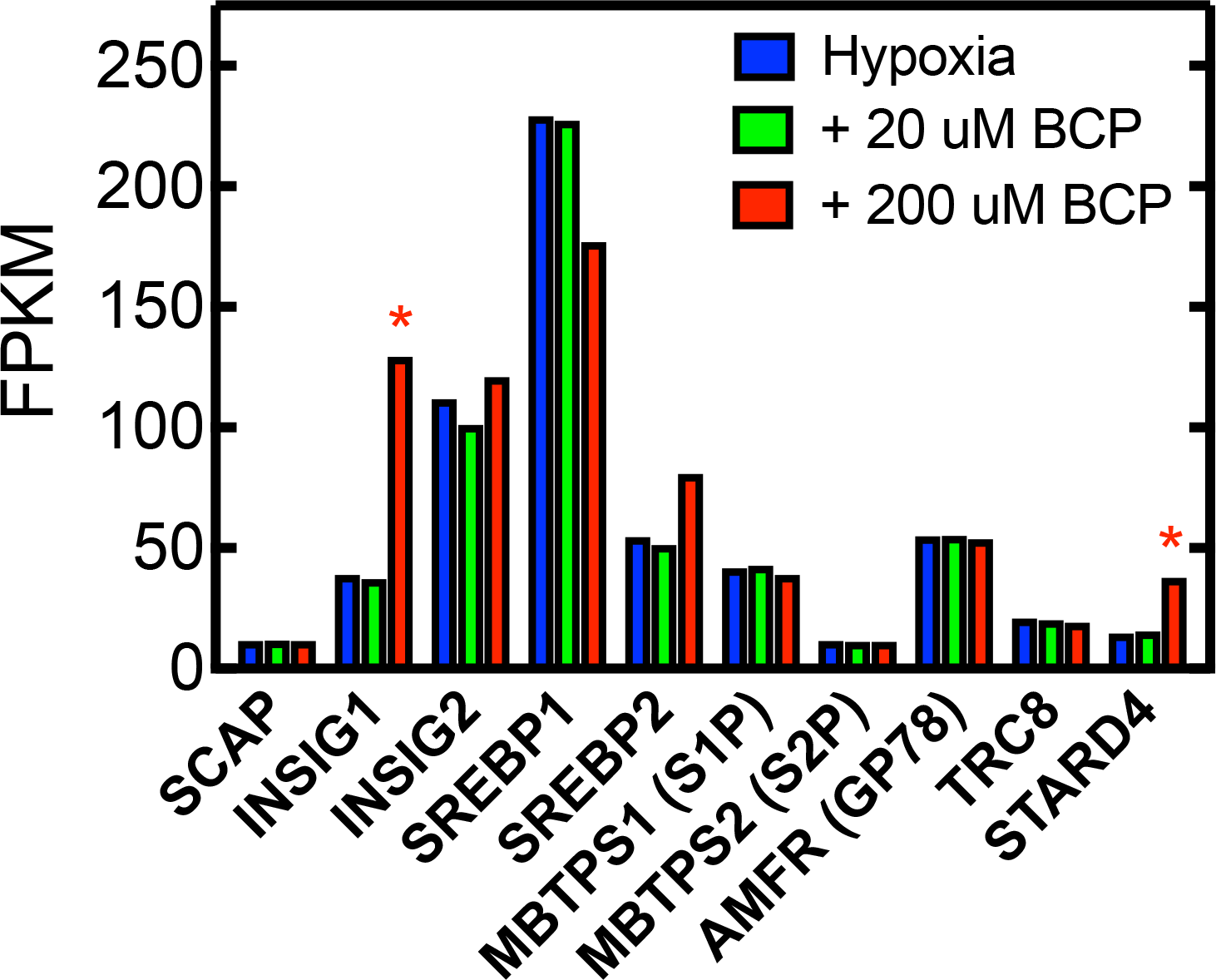
BCP induces transcription of INSIG1 and STARD4. Insig-1 binds to SCAP under cholesterol replete conditions, which prevents transport of SREBP isoforms to the Golgi. STARD4 is a cholesterol binding protein that shuttles cholesterol between the plasma membrane and the ER (and other membranes), and is under SREBP transcriptional control. * q ≤ 0.05.

### Simvastatin increases BCP-induced cytotoxicity

The action of BCP appears to be concentration dependent. At sub-cytotoxic levels is appears to promote lipid biosynthesis which may be used to membrane biosynthesis while a very high concentrations it induces cytotoxicity (cell death). We hypothesized that blocking cholesterol biosynthesis at sub-lethal concentrations of BCP would enhance cytotoxicity. To test this, we first analyzed the effect of two inhibitors (Atorvastatin and Simvastatin) of HMGCR on cell growth of UFH-001, T47D, and MCF10A cells using in the MTT assay (Figures 7A-C). The sensitivity of these drugs varied with cell type: UFH-001 > T47D > MCF10A based on IC_50_ values. In each cell line, Simvastatin was a better inhibitor of cell growth than Atorvastatin under our conditions. Simvastatin also sensitized UFH-001 cells to the cytotoxic effects of BCP (Figure 7), but this shift was only about 2-fold.

**Figure 7.**
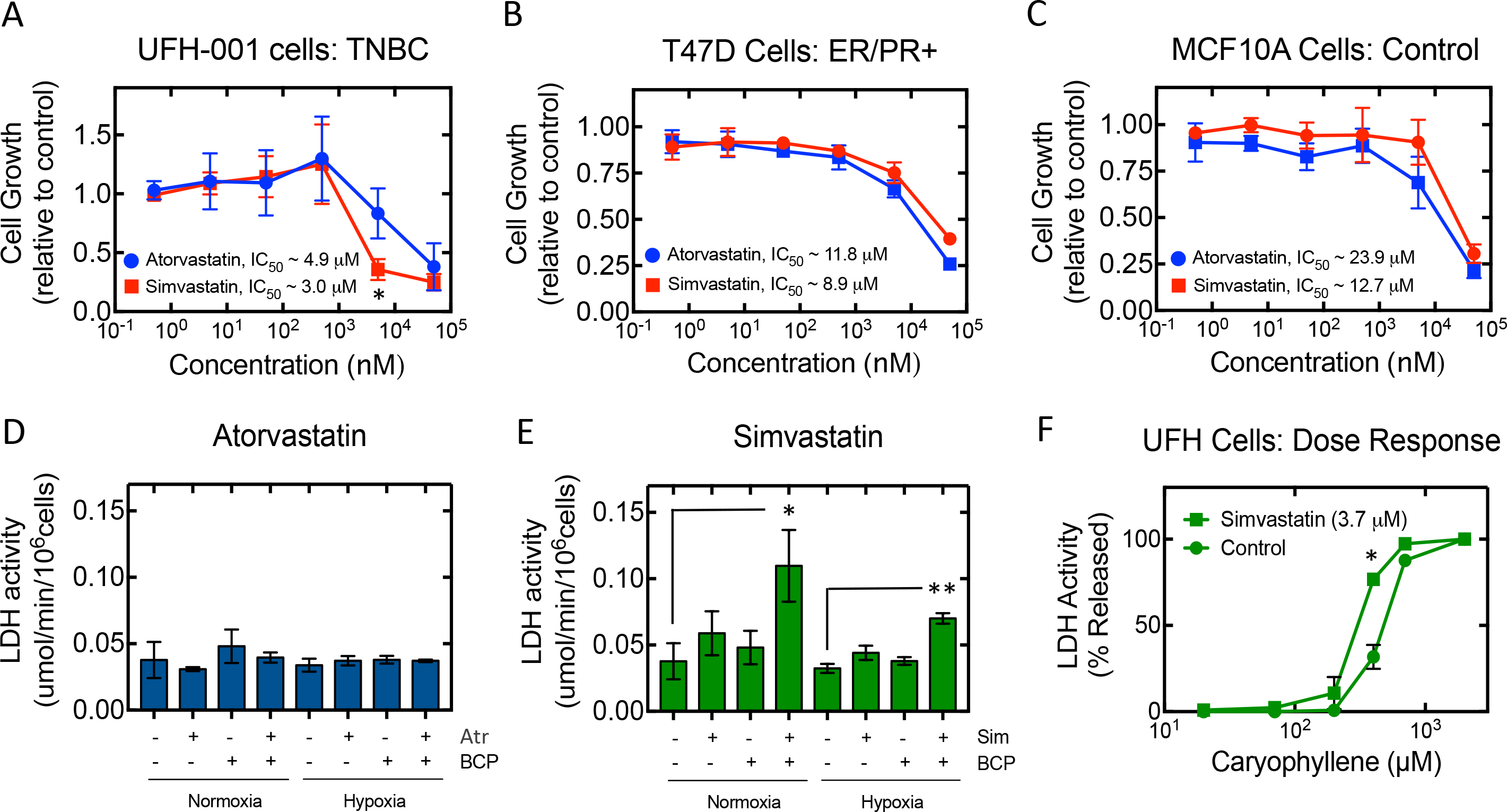
Simvastatin augments the cytotoxic effect of BCP in UFH-001 cells. **Panels A-C**. UFH-001, T47D, or MCF10 A cells were exposed to specific concentrations of Atorvastatin or Simvastatin for 48 h. The MTT assay was then used to assess viability/ cell growth. ID50 values for Atorvastatin and Simvastain are displayed in each graph. * p = 0.028. **Panels D and E**. UFH-001 cells were preincubated with Atorvastatin (D) or Simvastatin (E) for 32 h at their IC_50_ values. Then cells were then exposed (or not) to hypoxia in the absence or presence of BCP (200 μM) for 16 h. LDH activity was measure in the medium and reported as the μmol/min/106 cells (avg ± SEM). t-test was performed on Simvastatin samples. Normoxic samples treated with both Simvastatin and BCP vs Control: * p=0.046. Hypoxic samples treated with both Simvastatin and BCP vs control: ** p= 0.008. **Panel F**. UFH-001 cells were exposed to Simvastatin for 24 h, after which BCP was introduced for an additional 16 h. LDH activity was measured in the medium. Simvastatin induce a 2-fold shift in the cytotoxicity curve. * p = 0.016.

### BCP activates a feed back loop

Besides the genes that encode for the enzymatic reactions in cholesterol and lipid biosynthesis, we also evaluated the transcription of key regulators of cholesterol homeostasis (Figure 5). SREBP-1 transcripts were elevated relative to SREBP-2, which is the more selective over cholesterol biosynthesis. Although it would appear that transcription of SREBP-1 and SREBP-2 are regulated in opposite directions by BCP, neither of these changes achieved a q value of ≤ 0.05. When cholesterol levels are low, both SREBP-1 and SREBP-2 are escorted to the Golgi by the cholesterol sensing protein, SREBP cleavage-activating protein (SCAP). SCAP transcription is not affected by BCP. In the Golgi, the SREBPs encounter two specific proteases, S1P and S2P. These are responsible for the sequential cleavage of the SREBP’s to generate the “mature” transcription factor that then travels to the nucleus. Transcription of neither S1P nor S2P were enhanced by BCP. GP78 and TRC8 are E3 ubiquitin ligases that modulate the post-translation polyubiquitination of HMGCR to enhance cholesterol-dependent proteasomal degradation. Neither of these gene were regulated by BCP. SCAP localization depends on cholesterol levels in the ER. Above a certain threshold, SCAP is retained in the ER by the ER-resident proteins, insulin induced gene 1 or 2 (INSIG1 or INSIG2), which binds to SCAP preventing its chaperone function. The number of transcripts for INSIG2 was higher than that for INSIG1, but INSIG1 was significantly upregulated by BCP (q = 0.013). Finally, we measured the expression of STARD4 transcripts. STARD4 is a cholesterol specific transporter that moves cholesterol between membranes (39,40). BCP significantly stimulated transcription of this member of the START family member (q = 0.013). These later data suggest that BCP not only activates the biosynthetic path, but initiates a negative feedback system, as well, to ultimately turn off the biosynthetic process.

## Discussion

While BCP is cytotoxic to breast cancer cells at high concentrations, we show for the first time, that sub lethal concentrations induce the transcription of nearly all of the genes encoding proteins involved in cholesterol biosynthesis, and a significant few involved in fatty acid biosynthesis. This is reminiscent of the response caused by cholesterol depletion, where activation of the transcription factor, SREBP2, leads to the global upregulation of genes dedicated to cholesterol biosynthesis (41). There appears to be a very tight switch between the concentration of BCP that induces gene transcription and that leading to cytotoxicity. Because terpenes, in general, interact with membranes to induce swelling and changes in fluidity, it is possible that there is some specific level of membrane stress which is sensed by cells allowing them to mount an offense that attempts to replenish lost lipid, thereby maintaining membrane integrity. Indeed, others have shown that plasma membranes (PM) are not only targets of stress, but also sensors in activating stress responses (42). Any alterations in lipid fluidity within PM lipid microdomains, particularly that of the sphingomyelin and cholesterol rich environment of lipid rafts, leads to the re-organization of these domains followed by activation of specific signaling pathways (43). In cancer cells, changes in PM lipid composition, among other events, allows the recruitment of selective heat shock proteins like GRP78 (glucose-regulated protein 78) by the GRP78 co-factor, HJT-1 (44). GRP78 is normally a resident of the ER where it regulates protein processing and the unfolded protein response (45). Localization GRP78 to the PM of cancer cells exhibit therapeutic resistance (46). While transcription of GRP 78 (HSPA5) is not affected by BCP, we have not measured its translocation to the PM. However, we might expect this if alterations in PM lipid concentration, induced by BCP, serves to recruit GRP78 initiating the resistance response.

Is cholesterol content or redistribution within the PM the key to understanding BCP action? Cholesterol levels in membranes are tightly regulated. Ninety percent of the free cholesterol in cells is associated with the plasma membrane (PM), while smaller pools are found in intracellular membranes like endosomes, secretory vesicles, and caveolae (47,48). The ER, where the SREBPs and cholesterol sensing proteins reside, contains only about 1% of the cell’s cholesterol (47,49). This value represents a threshold in that above this concentration (i.e., 5 mol% of total ER membrane lipid), cholesterol blocks the activation of SREBP through interaction with its chaperone, SCAP (50). It is only recently that we have begun to appreciate the trafficking of cholesterol between the PM and the ER. Das et al. have shown that there are three cholesterol pools in plasma membranes (51,52). In their studies, a mutant form of a cholesterol binding protein (^125^I-labeled Perfringolysin O; ^125^I-PFO) was used to quantify cholesterol content in fibroblast PM. These authors discovered that one pool is accessible to ^125^I-PFO binding, which represents 16% of the PM lipids. This also represents the pool that signals cholesterol excess (or depletion) to the regulatory machinery in the ER. A second pool is associated with a compartment enriched in sphingomyelin. This pool is inaccessible to ^125^I-PFO, unless it is disrupted by treatment with sphingomyelinase. This pool may represent lipid rafts, the higher ordered PM domains that are enriched in sphingomyelin and cholesterol (53). On average, this pool represents 15% of PM lipids. A third pool, which was defined as the “essential” pool, accounts for about 12% of the PM pool and is not accessible, even after sphinomyelinase treatment, to ^125^I-PFO. This pool is required for membrane integrity. When cells are depleted of cholesterol, the PFO-accessible pool becomes depleted. Even thought the other two pools remain intact, there is no trafficking from these pools to the ER. However, when cholesterol-depleted cell are treated with sphingomyelinase, the liberated cholesterol restores the PFO-accessible pool, which is then able to communicate with the ER. It is possible that BCP reduces the concentration of the PFO-accessible pool in the PM leading to cholesterol depletion in the ER. It is also possible that the essential pool is affected which leads to loss of membrane integrity. It could be both, mediated by different concentrations of BCP. While we did not measure PM-specific pools, exposing radioactive cholesterol to BCP-treated cells increases the association of cholesterol relative to controls. As the PM is the major sink for free cholesterol, this infers that the PM pool is depleted by the action of BCP.

Blocking just 1% of cholesterol transport between the PM and ER leads to activation of cholesterol biosynthesis (54). Since cholesterol is essentially water insoluble, rapid nonvesicular transport will require carrier proteins. One interesting set of carrier proteins is the steroidogenic acute regulatory-related lipid-transfer (START) domain-containing family. The START domain binds cholesterol in a hydrophobic pocket covered by a “lid” that opens to allow exchange with membranes (39,40). One subgroup of this family contains only the START domain. Included in this group is STARD4 which increases the formation of cholesteryl esters in the ER (55). Mesmin et al. have shown that STARD4 can mediate transfer of cholesterol between membranes, and overexpression of STARD4 in cells increases the rate of transfer of sterol to the ER (56). Because the expression of STARD4 is under the control of SREBP-2 (57), this represents a negative feedback loop in which transport of cholesterol to the ER by STARD4 contributes the downregulation of cholesterol biosynthesis. Of note is the 2.5-fold increase in STARD4 transcription in the presence of BCP (q = 0.013). Thus, the mechanism of BCP appears complex in that it both activates the cholesterol biosynthetic pathway and at the same time induces a feed back loop to down-regulate that process. This is further evidence that the mechanism of BCP action is through cholesterol depletion because the same regulatory steps are affected by both BCP and cholesterol.

In contrast to the studies described herein, other investigators have reported hypolipidemic effects of BCP (58–61). These observations were detected, *in vivo*, under both hyperlipidemic and hypercholesterolemic conditions. In both models, the concentration of BCP that was achieved *in vivo* was about 130 μM. At this circulating concentration, BCP was able to reduce circulating cholesterol and LDL cholesterol in hyperlipidemic but not normal animals. The target of this action is inhibition of HMGCR activity. In addition, BCP (1.5 g/kg of diet, or 0.15%) inhibits tumor growth and lung metastasis of melanoma cells in a diet-induced obesity mouse model (62). In this model, BCP decreased lipid content in macrophage and adipocytes associated with the lung, and reduced the concentration of growth factors and cytokines associated with tumor tissue. They also showed that BCP suppressed VEGF expression, which is likely responsible for the observed reduction in tumor angiogenesis. Thus the action of BCP may depend on its concentration, the target tissue, and metabolic state of that tissue.

While simvastatin sensitized UFH-001 cells to the cytotoxic effects of BCP, this shift was only about 2-fold. However, this does open the possibility for combinatorial therapy if we could target cancer cells specifically. Preferential expression of GRP78 to the cell surface of tumor cells could serve that role. In fact, synthetic peptides mimicking GRP78 binding motifs fussed to cell death inducing peptides or cytotoxic drugs are able to promote apoptosis in cancer cells *in vitro*, including breast cancer cells (63). Another possibility is to change the structure of BCP to make it a better cytotoxic reagent. Terpenes are not the best starting material for drug development. However, studies from the Gertsch lab (64) showed that removing the conformational constraints induced by the medium-sized ring and by introducing functional groups at the sesquiterpene hydrocarbon 1 (see Figure 1 for carbon labeling), this new scaffold created a structure that not only binds to the CB_2_ receptor, but reversibly inhibits fatty acid amide hydrolase, the major endocannabinoid degrading enzyme. This might serve to prolong the effects of BCP, and make possible its use at lower concentrations.

In conclusion, we have shown that BCP induces a striking transcriptional upregulation of the cholesterol biosynthetic pathway, possibly providing cytoprotection for membrane structures and/or inducing drug resistance. However, blocking this path increases BCP-induced induced cytotoxicity suggesting a strategy for combinatorial therapy. In future studies, we will examine changes in membrane lipid and protein composition, the association with GRP78 translocation, and metabolic flux in lipid biosynthetic pathways in BCP-treated cells.

## Acknowledgements

The authors would like to recognize the exceptional cell culture skills of Xiao Wei Gu.

## Abbreviations

BCP: β-caryophyllene
SREBP: sterol response element binding protein
SCAP: SREBP cleavage activating protein
TNBC: triple negative beast cancer
SCD: stearoyl-CoA desaturase
HBrC: human breast cancer cells
LDH: lactate dehydrogenase
ER: endoplasmic reticulum
HMGCR: HMG CoA reductase
INSIG: insulin induced gene
S1P and S2P: site 1 protease and site 2 protease
STARD4: StAR related lipid transfer domain containing 4
GRP78: glucose-regulated protein 78
PFO: perfringolysin O
HER2: human epidermal growth factor 2

## References

1. International, W. C. R. F. (2012) Breast Cancer Statistics. in http://www.wcrf.org/int/cancer-facts-figures/data-specific-cancers/breast-cancer-statistics

2. Jones, S. (2008) Metastatic Breast Cancer: The Treatment Challenge. Clin. Breast Cancer 8, 224–233

3. Peddi, P. F., Ellis, M. J., and Ma, C. (2012) Molecular Basis of Triple Negative Breast Cancer and Implications for Therapy. Internat.J.Breast Canc. 2012, 7

4. Schneider, B. P., Winer, E. P., Foulkes, W. D., Garber, J., Perou, j. M., Richardson, A., Sledge, G. W., and Carey, L. A. (2008) Triple-Negative Breast Cancer: Risk Factors to Potential Targets. Clin.Cancer Res. 14, 8010–8018

5. Farmer, E. E. (2014) Leaf defence, OUP Oxford

6. Bayala, B., Bassole, I. H., Scifo, R., Gnoula, C., Morel, L., Lobaccaro, J. M., and Simpore, J. (2014) Anticancer activity of essential oils and their chemical components - a review. Am. J. Cancer Res. 4, 591–607

7. Gershenzon, J., and Dudareva, N. (2007) The function of terpene natural products in the natural world. Nature Chem. Biol. 3, 408–414

8. Hunter, W. N. (2007) The non-mevalonate pathway of isoprenoid precursor biosynthesis. J. Biol. Chem. 282, 21573–21577

9. Frost, C. J., Appel, H. M., Carlson, J. E., De Moraes, C. M., Mescher, M. C., and Schultz, J. C. (2007) Within-plant signalling via volatiles overcomes vascular constraints on systemic signalling and primes responses against herbivores. Ecol. Lett. 10, 490–498

10. Frost, C. J., Mescher, M. C., Dervinis, C., Davis, J. M., Carlson, J. E., and De Moraes, C. M. (2008) Priming defense genes and metabolites in hybrid poplar by the green leaf volatile cis-3-hexenyl acetate. New Phytologist 180, 722–734

11. Schiff, P. B. H., S.B. (1980) Taxol Stabilizes Microtubules in Mouse Fibroblast Cells. Proc. Natl. Acad. Sci. U S A 77, 1561–1565

12. Heinig, U., Scholz, S., and Jennewein, S. (2013) Getting to the bottom of Taxol biosynthesis by fungi. Fungal Diversity 60, 161–170

13. Perez, E. A. (1998) Paclitaxel in Breast Cancer. Oncologist 3, 373–389

14. Fidyt, K., Fiedorowicz, A., Strzadala, L., and Szumny, A. (2016) beta-caryophyllene and beta-caryophyllene oxide-natural compounds of anticancer and analgesic properties. Cancer Med. 5, 3007–3017

15. Dahham, S. S., Tabana, Y. M., Iqbal, M. A., Ahamed, M. B., Ezzat, M. O., Majid, A. S., and Majid, A. M. (2015) The Anticancer, Antioxidant and Antimicrobial Properties of the Sesquiterpene beta-Caryophyllene from the Essential Oil of Aquilaria crassna. Molecules 20, 11808–11829

16. Gertsch, J., Leonti, M., Raduner, S., Racz, I., Chen, J. Z., Xie, X. Q., Altmann, K. H., Karsak, M., and Zimmer, A. (2008) Beta-caryophyllene is a dietary cannabinoid. Proc. Natl. Acad. Sci. U S A 105, 9099–9104

17. Bayala, B., Bassole, I. H., Gnoula, C., Nebie, R., Yonli, A., Morel, L., Figueredo, G., Nikiema, J. B., Lobaccaro, J. M., and Simpore, J. (2014) Chemical composition, antioxidant, anti-inflammatory and anti-proliferative activities of essential oils of plants from Burkina Faso. PLoS One 9, e92122

18. Amiel, E., Ofir, R., Dudai, N., Soloway, E., Rabinsky, T., and Rachmilevitch, S. (2012) beta-Caryophyllene, a Compound Isolated from the Biblical Balm of Gilead (Commiphora gileadensis), Is a Selective Apoptosis Inducer for Tumor Cell Lines. Evid. Based Complement. Alternat. Med. 2012, 872394

19. Legault, J., and Pichette, A. (2007) Potentiating effect of beta-caryophyllene on anticancer activity of alpha-humulene, isocaryophyllene and paclitaxel. J. Phar. Pharmacol. 59, 1643–1647

20. Huang, M., Sanchez-Moreiras, A. M., Abel, C., Sohrabi, R., Lee, S., Gershenzon, J., and Tholl, D. (2012) The major volatile organic compound emitted from Arabidopsis thaliana flowers, the sesquiterpene (E)-beta-caryophyllene, is a defense against a bacterial pathogen. New. Phytol. 193, 997–1008

21. Tambe, Y. T., H.; Honda, G.; Ikeshiro, Y.; Tanaka, S. (1996) Gastric Cytoprotection of the Non-steroidal Anti-Inflammatory Sesquiterpene, β-Caryophyllene. Planta Med. 62, 469–470

22. Ghelardini, C. G., N.; Di Cesare, M.; Massanti, G.; Bartolini, A. (2001) Local Anesthetic Activity of β-Carophyllene. Farmaco 56, 387–389

23. Di Sotto, A., Evandri, M. G., and Mazzanti, G. (2008) Antimutagenic and mutagenic activities of some terpenes in the bacterial reverse mutation assay. Mutation Research/Genetic Toxicology and Environmental Mutagenesis 653, 130–133

24. Sarpietro, M. G. D., A.; Accollla, M. L.; Castelli, F. (2015) Interaction of Beta-caryophyllene and Beta-caryophyllene oxide with phospholipid bilayers: Differential scanning calorimerty study. Thermochimica Acta 600, 28–34

25. Sikkema, J., de Bont, J. A., and Poolman, B. (1994) Interactions of cyclic hydrocarbons with biological membranes. J. Biol. Chem. 269, 8022–8028

26. Holthuis, J. C., and Menon, A. K. (2014) Lipid landscapes and pipelines in membrane homeostasis. Nature 510, 48–57

27. Brown, M. S., Radhakrishnan, A., Goldstein, J. L. (2018) Retrospective on Cholesterol Homeostasis: The Central Role of Scap. Annu. Rev. Biochem. 87, 1.1–1.25

28. Chen, Z. A., L.; Mboge, M.Y.;McKenna, R.;Frost, C.J., Heldermon,C.D.; Frost, S.C. (2018) UFH-001 cells: A novel triple negative, CAIX positive, human breast cancer model system. Cancer Biol. Ther. 19, 598–608

29. Edgar, R., Domrachev, M., and Lash, A. E. (2002) Gene Expression Omnibus: NCBI gene expression and hybridization array data repository. Nucleic Acids Res 30, 207–210

30. Markwell, M. A. K., Haas, S. M., Lieber, L. L., and Tolbert, N. E. (1978) A Modification of the Lowry Procedure to Simplify Protein Determination in Membrane and Lipoprotein Samples. Anal. Biochem. 87, 206–210

31. Laemmli, U. K. (1970) Cleavage of Structural Proteins During the Assembly of the Head of Bacteriophage T4. Nature 227, 680–685

32. Li, Y., Tu, C., Wang, H., Silverman, D. N., and Frost, S. C. (2011) Catalysis and pH Control by Membrane-associated Carbonic Anhydrase IX in MDA-MB-231 Breast Cancer Cells. J. Biol. Chem. 286, 15789–15796

33. Chen, Z., Ai, L., Mboge, M. Y., Tu, C., McKenna, R., Brown, K. D., Heldermon, C. D., and Frost, S. C. (2018) Differential expression and function of CAIX and CAXII in breast cancer: A comparison between tumorgraft models and cells. PLoS One 13, e0199476

34. Keating, P., Cambrosio, A., Nelson, N. C., Mogoutov, A., and Cointet, J. P. (2013) Therapy’s Shadow: A Short History of the Study of Resistance to Cancer Chemotherapy. Front. Pharmacol. 4, 58

35. Li, Y., Wang, H., Oosterwijk, E., Tu, C., Shiverick, K. T., Silverman, D. N., and Frost, S. C. (2009) Expression and Activity of Carbonic Anhydrase IX Is Associated with Metabolic Dysfunction in MDA-MB-231 Breast Cancer Cells. Cancer Invest 27, 613–623

36. Horton, J. D., Goldstein, J. L., and Brown, M. S. (2002) SREBPs: transcriptional mediators of lipid homeostasis. Cold Spring Harb. Symp. Quant. Biol. 67, 491–498

37. Lanczky, A., Nagy, A., Bottai, G., Munkacsy, G., Szabo, A., Santarpia, L., and Gyorffy, B. (2016) miRpower: a web-tool to validate survival-associated miRNAs utilizing expression data from 2178 breast cancer patients. Breast Cancer Res. Treat. 160, 439–446

38. Goldstein, J. L., and Brown, M. S. (1990) Regulation of the mevalonate pathway. Nature 343, 425–430

39. Soccio, R. E., and Breslow, J. L. (2003) StAR-related lipid transfer (START) proteins: mediators of intracellular lipid metabolism. J. Biol. Chem. 278, 22183–22186

40. Alpy, F., and Tomasetto, C. (2005) Give lipids a START: the StAR-related lipid transfer (START) domain in mammals. J. Cell Sci. 118, 2791–2801

41. Brown, M. S., and Goldstein, J. L. (1997) The SREBP Pathway: Regulation of Cholesterol Metabolism by Proteolysis of a Membrane-bound Transcription Factor. Cell 89, 331–340

42. Vigh, L., Nakamoto, H., Landry, J., Gomez-Munoz, A., Harwood, J. L., and Horvath, I. (2007) Membrane regulation of the stress response from prokaryotic models to mammalian cells. Ann. N. Y. Acad. Sci. 1113, 40–51

43. Nagy, E., Balogi, Z., Gombos, I., Akerfelt, M., Bjorkbom, A., Balogh, G., Torok, Z., Maslyanko, A., Fiszer-Kierzkowska, A., Lisowska, K., Slotte, P. J., Sistonen, L., Horvath, I., and Vigh, L. (2007) Hyperfluidization-coupled membrane microdomain reorganization is linked to activation of the heat shock response in a murine melanoma cell line. Proc. Natl. Acad. Sci. U S A 104, 7945–7950

44. Birukova, A. A., Singleton, P. A., Gawlak, G., Tian, X., Mirzapoiazova, T., Mambetsariev, B., Dubrovskyi, O., Oskolkova, O. V., Bochkov, V. N., and Birukov, K. G. (2014) GRP78 is a novel receptor initiating a vascular barrier protective response to oxidized phospholipids. Mol. Biol. Cell 25, 2006–2016

45. Lee, A. S. (2014) Glucose-regulated proteins in cancer: molecular mechanisms and therapeutic potential. Nat. Rev. Cancer 14, 263–276

46. Zhang, Y., Tseng, C. C., Tsai, Y. L., Fu, X., Schiff, R., and Lee, A. S. (2013) Cancer cells resistant to therapy promote cell surface relocalization of GRP78 which complexes with PI3K and enhances PI(3,4,5)P3 production. PLoS One 8, e80071

47. Lange, Y. (1991) Disposition of intracellular cholesterol in human fibroblasts. J. Lipid Res. 32, 329–339

48. Lange Y., S., M. H., Ramos, B. V., Steck, T. L. (1989) Plasma Membranes Contain Half the Phospholipid and 90% of the Cholesterol and Sphingomyelin in Cultured Human Fibroblasts J. Biol. Chem. 264, 3786–3793

49. Sokolov, A., and Radhakrishnan, A. (2010) Accessibility of cholesterol in endoplasmic reticulum membranes and activation of SREBP-2 switch abruptly at a common cholesterol threshold. J. Biol. Chem. 285, 29480–29490

50. Radhakrishnan, A., Goldstein, J. L., McDonald, J. G., and Brown, M. S. (2008) Switch-like control of SREBP-2 transport triggered by small changes in ER cholesterol: a delicate balance. Cell Metab. 8, 512–521

51. Das, A. B., M. S., Anderson, D. D., Goldstein, J. L. Radhakrishnan, A. (2014) Three Pools of Plasma Membrane Cholesterol and their Relation to Cholesterol Homeostasis. eLife 3, e02882

52. Das, A., Goldstein, J. L., Anderson, D. D., Brown, M. S., and Radhakrishnan, A. (2013) Use of mutant 125I-perfringolysin O to probe transport and organization of cholesterol in membranes of animal cells. Proc. Natl. Acad. Sci. U S A 110, 10580–10585

53. Simons, K., and Sampaio, J. L. (2011) Membrane organization and lipid rafts. Cold Spring Harb. Perspect. Biol. 3, a004697

54. Infante, R. E., and Radhakrishnan, A. (2017) Continuous transport of a small fraction of plasma membrane cholesterol to endoplasmic reticulum regulates total cellular cholesterol. eLife 6, e25466

55. Rodriguez-Agudo, D., Ren, S., Wong, E., Marques, D., Redford, K., Gil, G., Hylemon, P., and Pandak, W. M. (2008) Intracellular cholesterol transporter StarD4 binds free cholesterol and increases cholesteryl ester formation. J. Lipid Res. 49, 1409–1419

56. Mesmin, B., Pipalia, N. H., Lund, F. W., Ramlall, T. F., Sokolov, A., Eliezer, D., and Maxfield, F. R. (2011) STARD4 abundance regulates sterol transport and sensing. Mol. Biol. Cell. 22, 4004–4015

57. Soccio, R. E., Adams, R. M., Maxwell, K. N., and Breslow, J. L. (2005) Differential gene regulation of StarD4 and StarD5 cholesterol transfer proteins. Activation of StarD4 by sterol regulatory element-binding protein-2 and StarD5 by endoplasmic reticulum stress. J. Biol. Chem. 280, 19410–19418

58. Tzeng, T. F., Lu, H. J., Liou, S. S., Chang, C. J., and Liu, I. M. (2014) Lipid-lowering effects of zerumbone, a natural cyclic sesquiterpene of Zingiber zerumbet Smith, in high-fat diet-induced hyperlipidemic hamsters. Food. Chem. Toxicol. 69, 132–139

59. Baldissera, M. D., Souza, C. F., Grando, T. H., Doleski, P. H., Boligon, A. A., Stefani, L. M., and Monteiro, S. G. (2017) Hypolipidemic effect of beta-caryophyllene to treat hyperlipidemic rats. Naunyn-Schmiedeberg’s Arch. Pharmacol. 390, 215–223

60. Baldissera, M. D., Souza, C. F., Doleski, P. H., Leal, D. B. R., Stefani, L. M., Boligon, A. A., and Monteiro, S. G. (2017) Enzymes that hydrolyze adenine nucleotides in a model of hypercholesterolemia induced by Triton WR-1339: protective effects of beta-caryophyllene. Mol. Cell. Biochem. 434, 127–134

61. Harb, A. A., Bustanji, Y. K., and Abdalla, S. S. (2018) Hypocholesterolemic effect of beta-caryophyllene in rats fed cholesterol and fat enriched diet. J. Clin. Biochem. Nutr. 62, 230–237

62. Jung, J. I., Kim, E. J., Kwon, G. T., Jung, Y. J., Park, T., Kim, Y., Yu, R., Choi, M. S., Chun, H. S., Kwon, S. H., Her, S., Lee, K. W., and Park, J. H. (2015) Beta-Caryophyllene potently inhibits solid tumor growth and lymph node metastasis of B16F10 melanoma cells in high-fat diet-induced obese C57BL/6N mice. Carcinogenesis 36, 1028–1039

63. Arap, M. A., Lahdenranta, J., Mintz, P. J., Hajitou, A., Sarkis, A. S., Arap, W., and Pasqualini, R. (2004) Cell surface expression of the stress response chaperone GRP78 enables tumor targeting by circulating ligands. Cancer Cell 6, 275–284

64. Chicca, A., Caprioglio, D., Minassi, A., Petrucci, V., Appendino, G., Taglialatela-Scafati, O., and Gertsch, J. (2014) Functionalization of beta-caryophyllene generates novel polypharmacology in the endocannabinoid system. ACS Chem. Biol. 9, 1499–1507

65. Tkachev, A. V. (1987) The Chemistry of Caryophyllene and Related Compounds. Khimiya Prirodnykh Soedinenii 4, 475–499

