## Supplemental Data for "β-Caryophyllene Enhances the Transcriptional Upregulation of Cholesterol Biosynthesis in Breast Cancer Cells"

Supplementary Figure Legends

**Supplementary Figure Legends**

**Supplementary Figure 1. Paclitaxel reduces growth of aggressive breast cells**. Panels A and B. UFH-001, MDA-MB-231 LM2, T47D, or MCF10 A cells were exposed to specific concentrations of Paclitaxel for 24 h (Panel A) or 48 (Panel B). The MTT assay was then used to assess viability/cell growth. Data are reported as percent of control: avg ± S.E.M., n = 3. Panel C. LDH activity was measured in the medium of cells treated with specific concentrations of Paclitaxel for 32 h followed by exposure to 70 µM BCP or vehicle for 16 h. Data are reported as the avg ± S.E.M., n=3. Panel D. Under the same conditions as in Panel C, the MTT assay was performed at 48 to assess viability/cell growth. Data are presented at OD at 570: avg ± S.E.M., n = 3.

**Supplementary Figure 2. Cytotoxicity of BCP is not affected by hypoxia**. Cells were exposed to normoxic or hypoxic condition in the presence of increasing concentrations of BCP. LDH release was measured after 16 h. Data shown represent the average of at least 3 biological replicates ± S.E.M.

Supplementary Figure 1

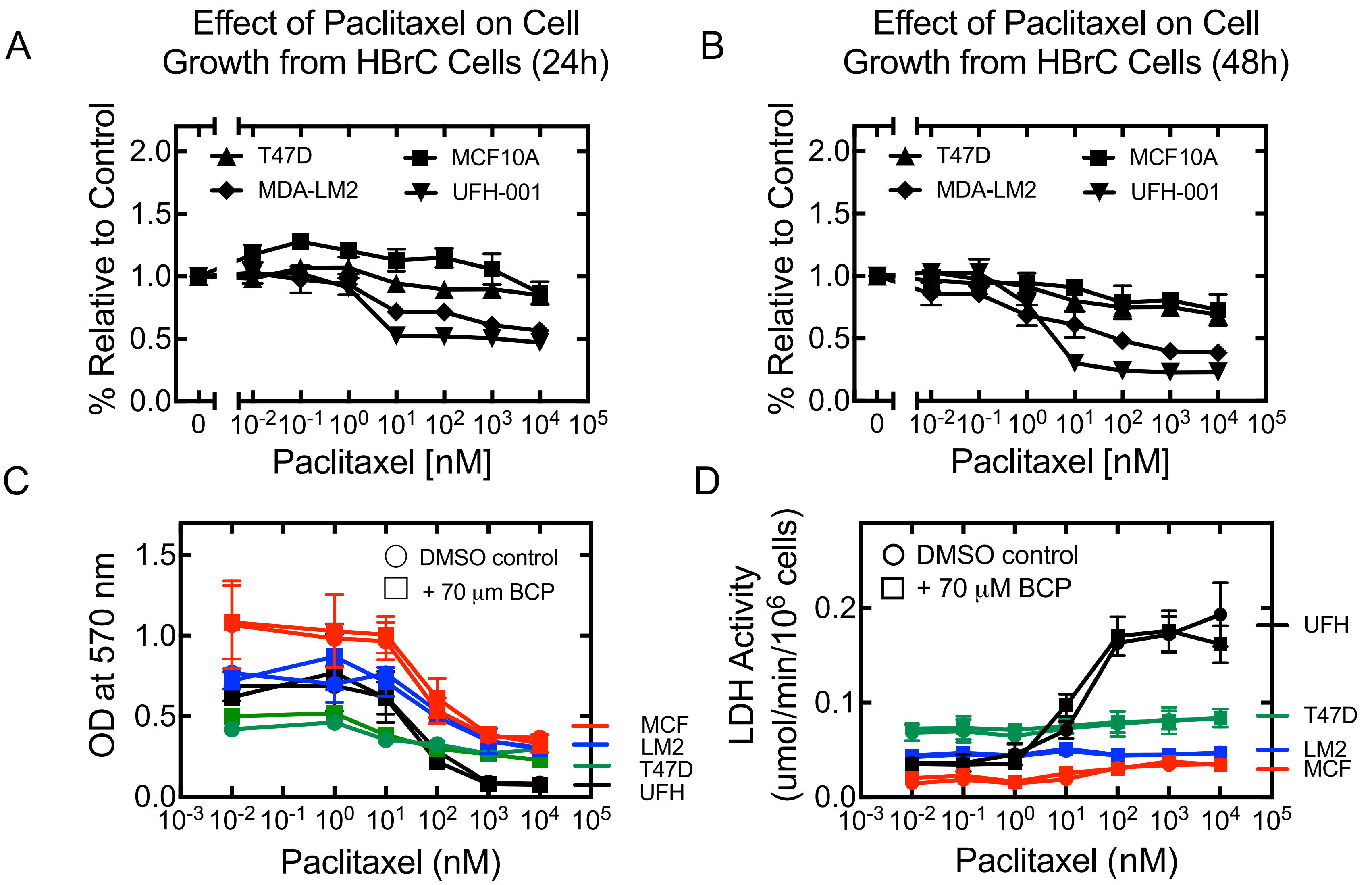

Supplementary Figure 2

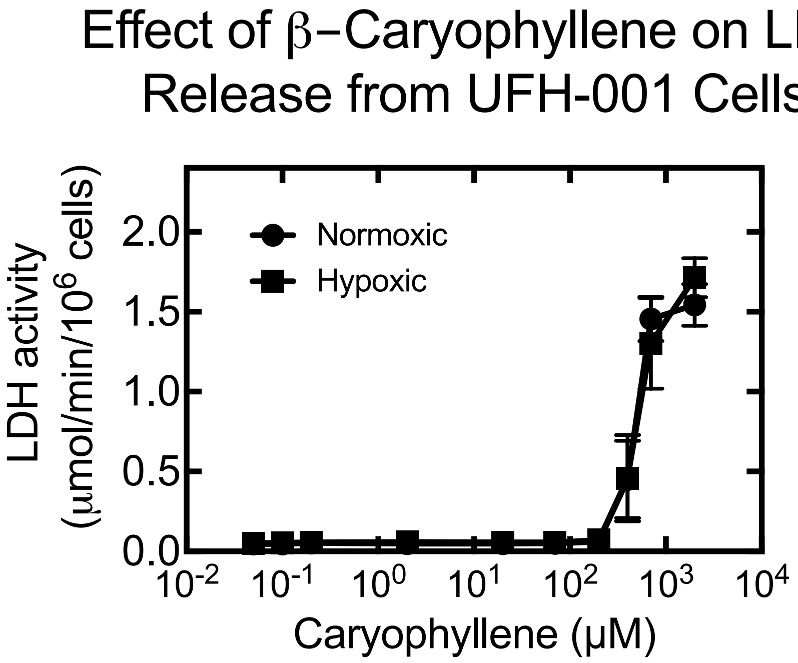
